# Maternal antibody-mediated elimination of a Puumala hantavirus outbreak in a bank vole colony

**DOI:** 10.1101/2025.11.04.686465

**Authors:** Stephan Drewes, Julia Wyszkowska, Ewa Jaromin, Joanna Hajduk, Ilona Onik, Mateusz Konczal, Krystyna Lach, Barbara Bober-Sowa, Katarzyna Baliga-Klimczyk, Edyta T. Sadowska, Rainer G. Ulrich, Paweł Koteja

## Abstract

Bank voles (*Myodes glareolus* syn. *Clethrionomys glareolus*) are frequently used as an animal model in ecological and biomedical studies, and are important reservoir of viral and bacterial zoonotic pathogens, e.g. of Puumala hantavirus (PUUV). Here we describe an accidental PUUV outbreak in a large bank vole laboratory colony by incursion of infected wild-trapped bank voles, and a successful eradication of the virus. The eradication plan was based on results of previous studies, which showed that maternal antibodies (MatAb) protect the young from infection for up to 40 days after the weaning, four weeks longer than the estimated duration of maintaining infectivity of PUUV in the environment. After ensuring that most animals are infected, 620 pairs were mated on the same day. Only females that showed PUUV-specific antibodies and gave offspring within 26 days after the mating were retained. All individuals of the parental generation were euthanized before the last weaning. The weaned offspring was moved to individually ventilated cages (IVC) and repeatedly tested for the presence of PUUV-specific antibodies and RNA. A few infected and suspicious animals were euthanised. Then the animals were mated (in IVC) and after producing grand-offspring euthanised and tested for PUUV RNA in lungs. No PUUV RNA was detected, and no animals showed PUUV-specific antibodies in next generations. The successful clearance confirmed the protective efficiency of PUUV-specific MatAb. The procedure for clearance of PUUV in the bank vole colony may represent a blueprint for similar approaches in precious colonies of other rodents infected by similar pathogens.

**Author Summary:** In 2006 we started a unique long-term experiment on the bank vole, a common European rodent. Our goal was to study how animals can adapt to different challenges - a process called adaptive radiation. We established 16 vole lines: four control lines and others selected for specific behavioural and physiological traits. Over the years, this colony became an important model for studying evolution, physiology, and behaviour. Unfortunately, the colony became infected with Puumala hantavirus. The virus is mild for voles but can cause serious zoonotic illness in humans, without specific medical treatment available. At first, it seemed that the entire colony would have to be destroyed - a loss of thousands of animals and many years of research. However, we used a natural advantage: young voles born to infected mothers are temporarily protected by maternal antibodies. By carefully planning breeding, isolation, and testing, we created conditions where the virus lost its strength before the young lost their protection. This simple yet challenging approach worked - we saved the colony. Because many animals respond to viruses in a similar way, our method can help rescue other valuable research populations without complex procedures like embryo transfer or cross-fostering.

## Introduction

Model animals characterized by unique traits are an invaluable and extensively used research tool both in basic and applied life sciences. Their great value has been typically paid by enormous time and financial investment. However, the laboratory colonies are endangered by infectious pathogens that may be difficult or seemingly impossible to treat, and hence may cause a drastic gene pool reduction or even loss of the entire colony. Even if the pathogens are not harmful to the host animals, veterinary or sanitary authorities can order elimination of the entire colony if the pathogen pose a hazard to people [1–4]. Therefore, efficient methods of eradicating such pathogens can be invaluable. Here we describe a successful eradication of Puumala hantavirus (PUUV; *species Orthohantavirus puumalaense*) from a unique colony of bank voles (*Myodes glareolus*, syn. *Clethrionomys glareolus*) comprising 16 genetically polymorphic lines of animals from a multidirectional selection experiment [5, 6].

The bank vole is a common rodent, widely used as a model species in ecological research (e.g., a classical monograph [7] or recent [8–12]) and in biomedical studies [13–15]. In the natural environment, the bank vole is the reservoir of several zoonotic pathogens, e.g. *Leptospira* spp. and PUUV [16, 17], but also most likely non-zoonotic agents, e.g. bank vole polyomavirus and bank vole hepaciviruses [18, 19].

PUUV belongs to the family *Hantaviridae* (order *Bunyavirales*) and shows a broad spatial distribution in Europe, reaching from northern and central Europe to the Balkans and Russia [16, 20–22] (more details are given in Discussion). PUUV transmission is exclusively horizontal and can be direct or via aerosolized excreta of infected voles. Viral RNA appears within 8-84 days post infection (dpi) in animals’ saliva, 11-44 dpi in feces and 14-44 dpi in urine; the peak level for these excreta is reached between 11-28 dpi [23]. Outside the host, PUUV remains infectious for up to 12-15 days at room temperature, but loses its infectivity faster at increased temperatures: at 37°C within 24h, and at 56°C within 15 min in wet cell culture, but a few hours at dry conditions [24]. Infected voles show PUUV-reactive antibodies in serum samples within 21 dpi or earlier [23, 25, 26]. Importantly, no vertical transmission from pregnant or lactating voles to their offspring was observed. This is because maternal antibodies (MatAb) transferred through placenta and milk temporally protect voles’ offspring against PUUV infection [27]. The authors showed that the offspring of infected mothers reared in an environment with PUUV did not develop their own antibodies until they were at least 80 days old. Thus, assuming that the time to develop antibodies is up to 20 days [23, 25, 26, 28], the offspring may be protected from infection up to about 60 days of age, but shorter if the maximum time to develop antibodies is in some individuals longer.

Humans can be infected with PUUV via direct contact with the infected vole (e.g. biting) or through inhalation of contaminated dust, but the virus is not transmitted between people. The infection can result in hemorrhagic fever with renal syndrome (HFRS), which severity is classified as mild to moderate in symptoms usually recalling a flu; however, 0.1-0.4% of fatal cases were reported [29]. Long-term monitoring studies indicated the association of increased numbers of human disease cases with bank vole reservoir population abundance and PUUV prevalence in reservoir populations [30–33].

Among pathogenic infections in laboratory rodents the viral ones are especially severe in consequences. In the case of a viral infection that poses a danger to humans, the respective veterinary or sanitary authorities typically enforce extermination of the whole colony, as was done in the case of an outbreak of lymphocytic choriomeningitis virus (LCMV) in a house mouse (*Mus musculus*) colony in USA [34]. This solution is well-founded considering that a) the intravital detection of infected individuals before they spread the virus is hardly possible (antibodies in blood appear too late for that purpose [35], b) the virus cannot be eliminated by any known pharmacological treatment, and c) although the immune system can often control the virus and the animals may not show obvious symptoms of disease, the animals chronically shed infectious virus, as is the case of bank voles infected with PUUV [35].

As the extermination of unique strains of laboratory animals is always a severe blow, alternative solutions are valuable. Because MatAb protect embryos, embryo transfer (ET) to non-infected surrogates could be considered as one such alternative. Although ET is feasible in laboratory mice, effective procedures for ET in other rodents, such as the bank vole, are not developed yet. Furthermore, ET is applicable when eradication of a virus from a few families is sufficient for rederivation of an inbred strain [36], but inefficient when two hundred or more families should be cleaned to preserve genetic heterogeneity in several genetically polymorphic lines. Another solution could be cross-fostering newborns to non-infected mothers [37–39], although the procedure is not always successful [38]. Cross-fostering of bank vole newborns is very effective [40], but again the efficiency of applying this procedure to a large population is doubtful, as another equally large non-infected laboratory colony should be available as a source of foster mothers. Moreover, transferring newborns from infected mothers carries a high risk of also transferring the virus, which would then infect healthy foster mothers.

Here we describe a successful direct eradication of PUUV from the 16 precious lines of bank voles [5, 6] without the need for embryonic transplantation or cross-fostering. The procedure was based on the assumption that MatAb would protect newborn and juvenile voles from infection for long enough to eliminate all PUUV sources before the voles became infected. Thus, this successful eradication confirms the high efficiency of MatAb in protection of offspring from PUUV infection. As the pattern of infectivity and MatAb protection is similar to that of other hantaviruses [37], we believe the same approach could have a much wider application.

## Results

### Detection of PUUV and identification of the virus origin

The colony was tested by ELISA for presence of PUUV-reactive antibodies in 2008, and the result was negative. In December 2012, however, serological analyses with two assays (in-house IgG ELISA and commercial Rapid test) performed on 17 old individuals (203 – 265 days of age) from generation 14 of the selection experiment showed that all of them were positive for PUUV-reactive antibodies (S1 Table). Subsequent S segment RT-PCR confirmed the infection for all 17 seropositive individuals. The pairwise nucleotide sequence similarities ranged between 99.1% and 100% (S2 Table). Similar pairwise nucleotide sequence similarities were observed for corresponding M and L segment sequences (S2 Table).

To identify the potential origin of the PUUV incursion, a phylogenetic analysis of the colony-derived PUUV sequences with reference sequences of PUUV strains of the various clades was performed and indicated a clustering of the colony-derived sequences with sequences from Teleśnica Oszwarowa, southern Poland, well-separated from sequences from Mikołajki, northern Poland and other sequences of the Russian clade (S1 Fig A-C). The pairwise comparison of partial S, M and L segment sequences showed a high similarity to sequences from Teleśnica Oszwarowa, southern Poland (S segment 98.0 – 99.8%; M segment 98.5 – 99.8%; L segment 99.2 – 99.7%), but much lower similarity to sequences from Mikołajki, northern Poland (S segment 82.8 – 83.5%; M segment 80.4 – 80.5%; L segment 86.6 – 86.8%) on nucleic acid level (S2 Table). These results indicated that the wild voles captured in Teleśnica Oszwarowa and temporarily held in the same facility were the source of the virus (see Materials and Methods).

Serological tests performed on humans after detecting the virus in voles showed that all those who worked directly with the infected bank voles and the technicians who were changing air filters in the ventilation system were anti-PUUV-antibody-positive. Importantly, however, all five tested bank voles representing another colony maintained in the same facility were not infected, even though they were maintained in rooms not separated by a sanitary barrier from those of the infected colony (S1 Table). Similarly, people who worked only with voles from the other colony or worked in the facility without direct contact with the voles, as well as people who worked in the same building but outside the facility, were not positive for PUUV-specific antibodies. The results implied that the temporary maintenance of the colony did not pose an immediate risk to people who had already worked with the infected colony and had developed immunity [41], or to people working in the same building but without contact with the infected voles or contaminated dust. This was the basis for the decision of sanitary authorities to not exterminate the colony immediately, but give us some time for the attempted eradication of the virus (initially 6 months, prolonged then to 10 months).

### Outline of the eradication program

The plan of PUUV eradication was founded on two already mentioned reports: (a) outside the host organism, PUUV loses its infectivity at room temperature within a maximum of about 15 days [24], but (b) offspring of infected mothers are protected by MatAb, presumably at least until 60 days of age [27]. The voles in our colony are weaned at the age of 17 days. Thus, the virus transferred from the maternal cage on the bodies/fur of juveniles is likely to remain infectious only until the voles are 32 days old (17 plus 15 days). This gives, optimistically, a “safety time window” of 28 days (60 minus 32 days) or longer before protection by MatAb becomes inefficient. Therefore, theoretically, if all the voles from the parental generation (genP) are reproduced at the same time and euthanized immediately after weaning their offspring on day 17, the virus should be not effectively transmitted to the offspring generation (genO).

This apparently simple plan, however, was based on assumptions that could have been overly optimistic. Moreover, it posed several technical problems, particularly challenging when one considers that, in order to preserve the genetic variation in the 16 lines, we aimed to eradicate the virus from the offspring of at least 240 families, i.e. more than 1000 individuals, in a facility that did not guarantee full protection against virus transfer between cages. We will address all these issues in Discussion.

The major problem was that the realistic plan had to assume some offspring would be not fully protected by MatAb and would become infected. Therefore, the offspring had to be kept in individually ventilated cages (IVC), ideally of the type warranting full protection against virus transmission, which was far beyond the financial reach. Instead, the weaned voles were placed in AERO Mouse Green Line IVC (Tecniplast, Bugugiatte, Italy), which, although equipped with HEPA filters, were not designed to warrant full protection against virus transmission. Next, the animals and their environment had to be frequently tested to detect and eliminate infected individuals before they infect the others. However, negative tests for the presence of antibodies or the viral RNA in samples that could be taken *intra vitam* (faeces, blood) do not provide a firm evidence of the virus absence. The ultimate negative evidence can be obtained only by *post mortem* analyses. Therefore, to the simple “one-step” plan outlined above we added another security step. After checking that none of the adult genO voles were seropositive, they were mated to produce another generation (grand-offspring: genG). After weaning the genG, all the genO individuals were euthanized and tested for the presence of PUUV RNA in lung samples.

### The eradication programme application

#### Procedures on parental generation (genP)

The eradication programme started on 1 February 2013 (day 1; Fig 1). Preliminary tests for the presence of PUUV-reactive antibodies (Ab-Dect test) performed on 45 females at the onset of the programme indicated that none of 12 individuals aged 94-117 days was seropositive, 8 of 12 individuals aged 121-131 were seropositive, and all individuals older than 131 days were seropositive (Fig 2). The result indicated that MatAb can provide the protection at least as long as reported in the previous studies, but also that the reproduction should be postponed until all potential mothers are seropositive. It is also worth noting, that above the age of 131 days, the level of antibodies was not clearly correlated with age, and that among the oldest individuals both very high and low values were observed (just above the “positive” threshold of 15; Fig 2).

**Fig 1.**
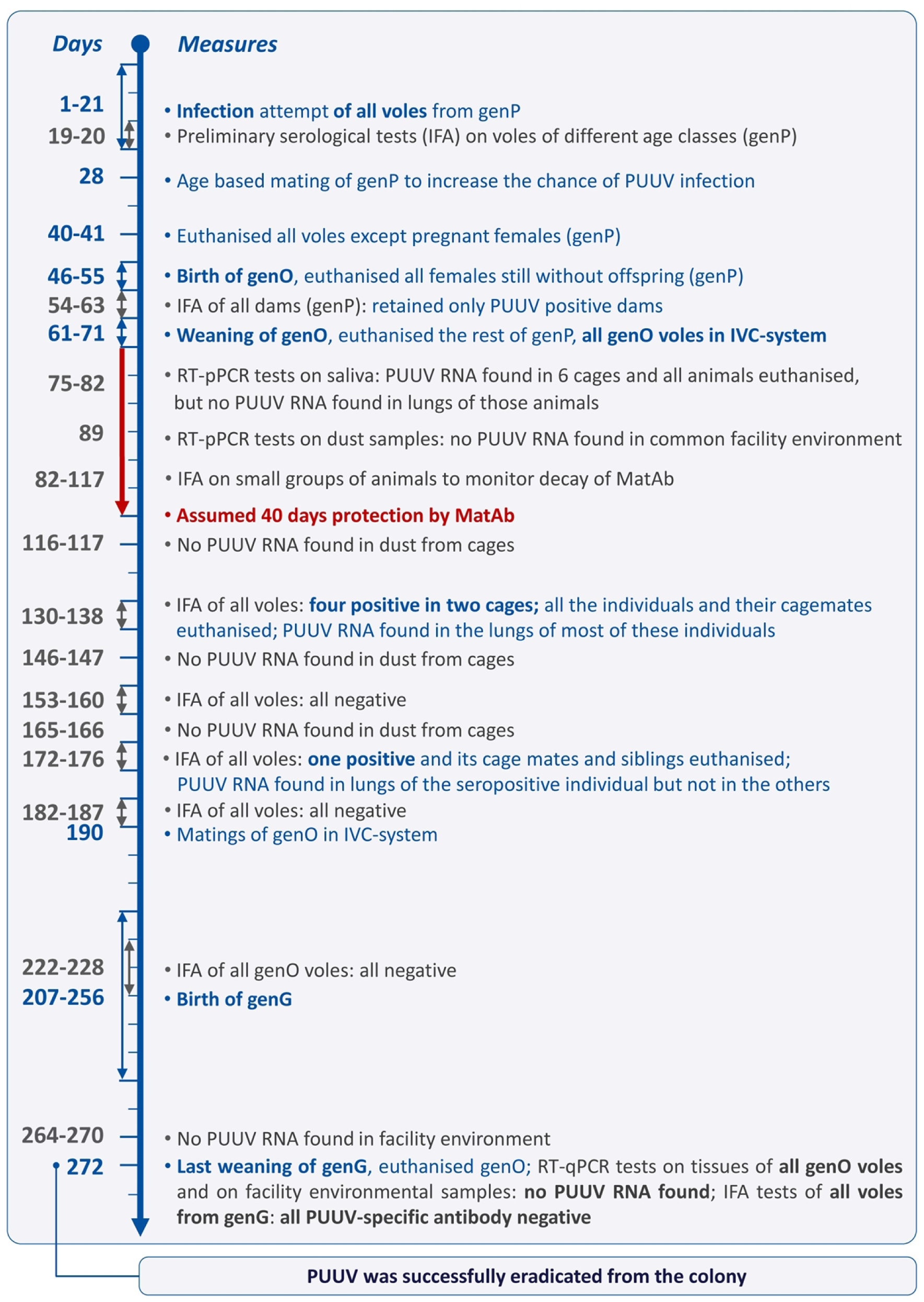
Puumala virus eradication scheme reflecting taken measures per day. Virus eradication measures are given in blue and diagnostics for Puumala virus surveillance in grey. Assumed days of protection according to [27]. Abbreviations: genP: parental generation (selection experiment gen.=15), genO: offspring generation (gen.=16), genG: grand-offspring generation (gen.=17), IVC: individually ventilated cages, IFA: indirect fluorescent antibody assay, MatAb: maternal antibodies, RT-qPCR: quantitative reverse transcription-polymerase chain reaction, PUUV: Puumala virus.

**Fig 2.**
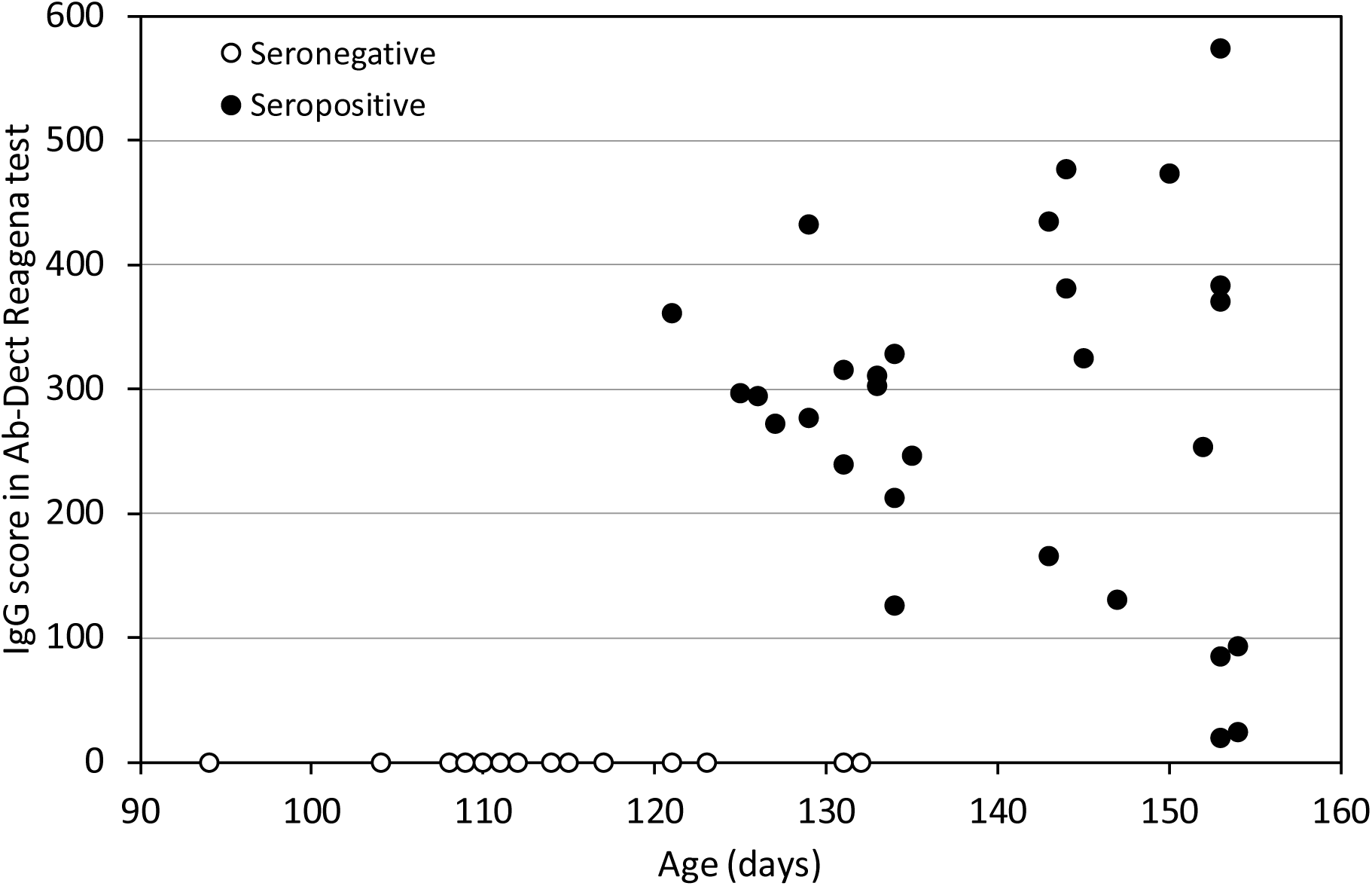
The relation between anti-PUUV IgG antibody level measured as a score in Ab-Dect Reagena test and age. The results were obtained for 45 females from the parental generation (genP) tested to assess the age at which most animals are seropositive. The threshold values are: negative < 5, ambiguous (not observed), positive > 15.

Preliminary tests for the presence of PUUV-reactive antibodies (Ab-Dect test) performed on 45 females at the onset of the programme indicated that none of 12 individuals aged 94-117 days was seropositive, 8 of 12 individuals aged 121-131 were seropositive, and all individuals older than 131 days were seropositive (Fig 2). The result indicated that MatAb can provide the protection at least as long as reported in the previous studies, but also that the reproduction should be postponed until all potential mothers are seropositive. It is also worth noting, that above the age of 131 days, the level of antibodies was not clearly correlated with age, and that among the oldest individuals both very high and low values were observed (just above the “positive” threshold of 15; Fig 2).

To ensure that all females are infected, used bedding was mixed between cages (4, 3 and 2 weeks before the planned mating). Another series of preliminary tests, performed on days 19-20 with the indirect fluorescent antibody assay (IFA), showed that 12 of 24 individuals aged 108-119 days were seropositive, and only 9 of 25 individuals aged 121-126 days were positive. Thus, the results of both preliminary tests indicated that, for the purposes of the eradication programme, females should be mated at an age of at least 130 days. Furthermore, all females giving birth must tested for the presence of anti-PUUV antibodies before their offspring are included in the next steps of the programme.

On day 28, 620 pairs were mated (35-40 per each of the 16 lines) and all remaining voles were euthanized. To increase the chance that only immunized females are mated, and also to increase the reproductive success, wherever possible older individuals were chosen (130-181 days old), but to ensure adequate representation in each of the lines, 25 females aged 117-128 days were also included. Wherever possible, the regular selection criteria used in the selection experiment were considered, too [5]. Males were euthanized 12-13 days after the mating (day 40-41). From day 17 after mating females were daily checked for pregnancy and birth, and the non-pregnant females were euthanized. The first births occurred 18 days post mating and within 11 days 473 females gave birth (mostly in the first 6 days), and the remaining females were euthanized. On day 7-12 after parturition, cages were changed and IFA tests were performed on all females that gave birth. Further, we retained only mothers that a) were seropositive, b) gave birth up to 26 days post mating (to narrow the age span in the cohort of offspring to 9 days), and c) had the litters of 3 or more young. From those that met these criteria we choose 280 (17-18 per line), whose offspring were weaned to IVC (days 63-71). Immediately after the weaning, all genP voles and excess offspring were euthanized.

Successively with reduction of the colony all the animal rooms and the common work areas in the animal facility were carefully disinfected, including the ventilation system. Since day 71 of the eradication programme all potential infectious material was confined in the IVC system, placed in one, previously disinfected room, in which all further work with the animals was performed. Tests for the presence of PUUV RNA in dust sampled from several places of the facility, performed repeatedly later on, gave negative results, which showed efficiency of both the disinfection and isolation procedures.

#### Procedures on offspring generation (genO)

The whole weaned litters were immediately placed in individually ventilated cages (IVC) equipped with HEPA filters (model G500, Aero “Green Line” system, Tecniplast, Bugugiatte, Italy), operating at 10% under-pressure and air flow rate of 75 volume exchanges per hour. As the priority was to preserve genetic pool, and hence to keep the offspring from as many families as possible, litters were reduced to 6 pups, so that each family could occupy only one of the 280 available IVC.

In an attempt to reduce the duration of PUUV infectivity in the environment, the room temperature was set to 26°C instead of the usual 20°C for the first two weeks, until the end of the first round of cage and bedding changes. To prevent within-sibling mating, the litters had to be sexed before the voles reached sexual maturity. However, because of the limited number of IVC, same-sex individuals from different families must have been joined (up to six individuals from two families). This increased the risk of transmitting the virus from an infected family to others, and also increased the risk of losing animals, especially males, as a result of inter-individual aggression. Therefore, to delay sexual maturation, already after the weaning the photoperiod was changed for short day (9L:13D). Separation of sexes was performed at the age of 28-39 days (days 84-85 of the eradication programme).

In an attempt to early infection detection, we took saliva samples for PUUV RNA analyses from 240 cages (pooled samples from all animals in a cage) 11-12 days after weaning (days 75-82). Viral RNA was detected in samples from six cages, and all 30 individuals from these cages were euthanized. However, a subsequent analysis of RNA extracted from their lungs indicated that the animals were not infected. Thus, the positive signal was probably due to contamination of the saliva sample with a dust from fur, which could still contain the virus (or inactive viral RNA) transferred from maternal cages. Concerning the difficulty to avoid such false positive signals, we resigned from further attempts to analyse saliva samples. After two cycles of cage changes, on days 116-117 dust samples from IVC were taken for the RNA analyses. Later, samples of the dust from cages and environment were then analysed repeatedly. All the results were negative, with a few exceptions described below.

Between day 82 and 117, i.e. at the age of 36 to 71 days, 6 series of IFA tests were performed on small groups of animals (9, 17, 26, 14, 15 and 24 individuals, partly repeated) to monitor the decay of PUUV-specific MatAb. Among 17 individuals 36-42 days old, MatAb were present in 13 individuals, absent in one, and in three the result was uncertain. At the age of 49-50 days, 12 individuals were positive, 9 negative, and 4 uncertain, and at the age of 56-64 only 3 were positive, 12 were negative and 14 uncertain. Of 24 individuals tested at the age of 71 days (all the first time), only 1 individual was positive, 16 were negative and 5 uncertain. Therefore, ‘positive’ results from tests performed at a later age should be considered to be highly indicative of PUUV infection.

On days 130-139, i.e., at the age of 76 days, the IFA tests aimed at detection infected voles were performed on all individuals. The test was positive for three individuals (and ambiguous in two more) from two cages (representing three families). All the individuals and their cage mates were euthanized on day 134. Analyses of lung tissue confirmed the presence of viral RNA in all the seropositive individuals, as well as in two of their four cage mates. Viral RNA was found also in dust taken from these cages. However, the IFA tests were negative in their sibling maintained in separate cages. The results were still negative in three subsequent tests, also in a test performed on day 166, i.e., 82 days after separating the sibling. However, a next test, performed on day 175, revealed a positive result in a single individual, a sibling of one of the infected individuals detected earlier (euthanized on day 135). Therefore, the single sibling of the newly detected individual (who was kept in the same cage) and the remaining siblings of the previously infected animals (who were kept in separate cages) were euthanized as well, even though they were PUUV-negative. The test for presence of PUUV RNA in lungs showed that the seropositive individual was indeed infected, but all the other individuals were not.

All tests for the presence of PUUV RNA in the facility environment were negative, except for a dust sample taken on day 160 from the exhaust air collector of the IVC system segment of 140 cages, among which there were the two with the infected animals eliminated on day 134. However, the analyses could not determine whether infectious PUUV or only its RNA remains were detected in the analysis performed 26 days after the last infected animal was removed.

#### Rearing the grand-offspring generation (genG)

A next series of IFA tests, performed on days 182 – 187, revealed no seropositive individuals, and no viral RNA was detected in dust sampled from cages. However, the previous observation of the single infected individual, in which the antibodies were still not detectable 82 days after the last contact with an infected individual, but were detected on day 92 after the last contact, showed that the negative result of the IFA test could not be regarded as proof that the voles do not carry the virus. The tests for the presence of viral RNA in lungs can provide such evidence, but require sacrificing the animals. Therefore, the next, grand-offspring (genG) generation had to be produced under the controlled conditions, before the eradication could be confirmed.

Therefore, photoperiod was reset to long day (16L: 8D) and the voles were again mated on day 190. As the entire procedure had to be performed in the IVC system, the number of pairs was limited to 280 (the breeders were chosen to maximize representation of independent families within each of the lines). All the genO voles were again tested for the presence of PUUV-specific antibodies, and the results were negative. All reproductive females were euthanized after weaning the offspring, and all other genO individuals were euthanized earlier. No PUUV RNA was found in their lungs nor in the environment of the animal rooms.

Thus, as voles from the “offspring” generation were not infected, and the environment was also PUUV free, young voles from the “grand-offspring” generation could be declared PUUV-free, too. The successful eradication was then additionally confirmed by negative IFA tests for PUUV-specific antibodies and RNA presence in the subsequent two generations.

## Discussion

### Identification of the PUUV origin in the bank vole colony

Molecular investigations suggest local evolution of PUUV strains, but might suggest also migration and invasion processes post vole population collapses [42, 43]. The current distribution of PUUV lineages is mainly shaped by the postglacial immigration of bank vole phylogroups [44]. Thus, central and western Europe was invaded by the western phylogroup of the bank vole distributing the central European PUUV lineage [45]. In contrast, in eastern and northern Europe the situation is much more complex. Different PUUV lineages can be harboured by different phylogroups of the bank vole. So far, in northern Poland exclusively the Latvian PUUV lineage was detected in bank voles of the Carpathian and Eastern phylogroups, whereas in southern Poland PUUV of the Russian lineage could be demonstrated to be harboured by bank voles of the Carpathian and Eastern phylogroups [46, 47].

PUUV sequences of the 17 bank voles from generation 14 were almost identical suggesting a single incursion event. To identify the potential origin of the PUUV incursion, these colony-derived sequences were compared to reference sequences of PUUV strains of the various clades indicating a high nucleotide sequence similarity and phylogenetic clustering with sequences from Teleśnica Oszwarowa, southern Poland. Therefore, the infection probably originated from the group of voles that were trapped in Teleśnica Oszwarowa and temporarily maintained (2009–2011) in the same facility as the selection experiment colony. Based on the eradication experiment described here, future analyses will allow to determine the mutation rate of PUUV in bank voles and its comparison to the situation in cell culture [42].

### The eradication scheme and its pitfalls

The simple plan for PUUV eradication outlined in the second section of Results was based on two assumptions supported by fairly convincing empirical data: (a) that PUUV loses its infectivity outside of the host at room temperature within about 15 days [24], and (b) that the offspring of infected mothers is protected by MatAb until the age of 60 days or longer [27], and on a simple calculation leading to the conclusion that if the young voles are weaned and separated from any infected mother at the age of 17 days, the virus transmitted on their bodies will become non-infectious approximately 4 weeks before protection by MatAb ceases.

Nevertheless, the plan was vulnerable to many hitches, which should be considered by other researchers who would need to adopt this scheme in their work:

1. The estimates of both the MatAb protection period and the PUUV infectivity duration in environment were based on small or only moderately sized samples. Thus, neither the minimum of the former nor the maximum of the latter were established with high reliability. If the actual extremes deviated from the reported values by just 10 days, the safety window would become very narrow.
2. The safety window considered for the entire colony is in practice much shorter because, even if all parents are mated at the same time, the parturitions are spread over several days, which again narrows the safety window.
3. The plan requires that all mothers are infected a long time before mating and have developed a sufficiently high level of PUUV-specific antibodies to provide their young with effective protection by MatAb. To help achieve this, we used a paradoxical approach of intentionally infecting all animals before implementing the proper eradication procedure. However, the available quantitative tests for the antibodies require large amount of blood, and practically could not be performed without euthanizing the animals. On the other hand, the available rapid immunoassays need only a drop of blood, but provide only qualitative or semi-quantitative results. To our knowledge, no studies were performed to find out whether and to what extent the efficiency of the MatAb-mediated protection changes with the level of anti-PUUV antibodies in mother. Thus, a mother “positive” according to such a test could still not provide effective protection to the offspring.
4. Importantly, even a single non-completely protected juvenile would cause re-occurrence of the infection in the next generation, unless the individual is isolated and eliminated before transmitting the virus. However, the virus starts spreading before PUUV-specific antibodies in blood can be detected [23]. Moreover, the simple assays do not allow to distinguish whether a young individual still carries MatAb or started to produce its own as a result of infection. Tests for the presence of viral RNA in saliva or excreta may provide a positive evidence of infection, but do not provide a negative evidence (that the animal is not infected), because the virus shedding is discontinuous [23]. Samples from the cage environment could be a more effective measure, but if applied soon after the weaning they would result in false positives, because some amount of virus or just viral RNA may be carried from maternal cages on the juveniles’ bodies. To summarize, no *intra vitam* method allows a reliable detection of the infection before the virus starts spreading, and actually not even at the time when the spreading already occurs.
5. Thus, the plan must assume that some of the young individuals could be infected. Therefore, all the young animals must be kept in isolation, which requires application of IVC and appropriate cage-changing station to prevent the virus transmission during animal maintenance procedures and blood testing. Such an equipment, of the protection level designed for preventing viral transmission (grade III), and the capacity sufficient to maintain several hundred individuals, was far beyond our financial reach, and the same may be true for most other potentially interested researchers. We could afford to purchase only the IVC system of grade II+ (“Aero Green line”, Tecniplast, Italy), which – although equipped with HEPA filters – is not warranted to prevent viral transmission. It did work good enough – the few infected animals found in two cages did not spread the virus to others. However, we cannot be sure whether the protection can be indeed regarded as reliable, or we were just lucky.
6. We initially assumed that the absence of PUUV-specific antibodies after a 55-day quarantine (a maximum 15 days of PUUV infectivity outside the host plus a maximum of 40 days for developing antibodies) can be treated as an evidence of successful eradication. However, the case of the single vole that did not show antibodies in the test used 82 days after the last contact with an infected vole or infectious material, but which has been detected as seropositive by day 92, changed the perspectives radically. In practice, we have decided that no period can be assumed to be long enough to provide the negative evidence, and therefore the one-step eradication plan has no *a priori* defined moment at which success or failure can be definitively declared.
7. The negative evidence can be provided only by *post mortem* tests for the presence of the viral RNA in lungs (or perhaps other organs), and also tests for the presence of the viral RNA in the environment, which have proved to be extremely sensitive, and allowed the detection for at least a month after the infectious virus material was removed. Therefore, the eradication programme must *a priori* plan the second step: producing within the IVC system a next generation of offspring, and performing the RNA analyses in lungs of all parents (or in another organ in which a particular virus is residing).
8. This raises the question of whether it would not be better to schedule the second reproduction as early as possible, i.e., just after the animals reach sexual maturity. In the case of bank voles, this would be at around 60 days of age, which would considerably shorten the whole procedure. However, we would not recommend this, because of increased risk that some of the individuals are infected even though not yet detected as anti-PUUV-positive by the serological test, and hence an increased risk of transmitting the virus to the grand-offspring generation.
9. This was not a planned experiment, but rather a ‘rescue operation’ carried out on a tight budget and with ‘fast-track design’, i.e. instant decisions about the next steps based on current information. Consequently, not all procedures were well optimised and some decisions were imprudent.
10. Therefore, if we could, we would introduce to the procedure a few changes: First, it would be better to test all the lactating females with the more reliable, semi-quantitative test (Ab-Dect Reagena), rather than qualitative IFA test. The variation of the level of antibodies was presumably very high (Fig 2), and the tests would allow to select for further steps only offspring of females that had high level of antibodies. Similarly, all the weaned young could be individually tested with the semi-quantitative test for the level of MatAb, and only those with high level retained. Possibly, in this way we would avoid having non-completely protected young in the offspring generation, and the entire procedure would be shortened. We could not financially afford such tests (roughly 10 times more expensive than the IFA tests), but in the case of attempted eradication of the virus from a smaller population this should be certainly considered as a plausible improvement. Second, we would not attempt to detect viral RNA in the saliva of young animals 12 days after weaning. This is because contamination of the samples with the virus transmitted at weaning, which is still present in the environment, leads to “false alarms” and the unnecessary euthanising of animals. In fact, it is not worth to invest in the tedious attempts to sample saliva, which inevitably increases also the chance of transmitting the virus between cages. On the other hand, sampling dust from the cages is easy, provides an integrated information about the virus shedding over several days, and the tests on dust have proven to be highly sensitive. Again, however, such tests should only be performed after a longer period since weaning and after at least two rounds of cage changes. Third, after detecting the first few infected individuals about 50 days after the weaning, we eliminated only their cage mates, but not their sibling and cage mates of their sibling, which were not seropositive. We did this because, at that time, we did not know how many of the young animals were infected. After losing many animals due to a ‘false alarm’, we were reluctant to eliminate more without evidence of infection. Instead, the suspicious animals were tested more frequently. However, from the perspective of further experience – detecting the infected sibling more than 40 days later – we see that this decision was wrong, and that all those suspicious animals should have been eliminated, too.

## Conclusion

In the presented case, we demonstrated that, once the pattern of pathogen infectivity is understood, it is possible to eradicate viral contamination from an animal facility by implementing improved logistics and moderately advanced safety precautions. This approach, enables the valuable colony to be maintained without drastically reducing the population size (compared to other methods such as embryo transfer or cross-fostering), thus preserving its genetic polymorphism. Given the scientific and financial benefits of maintaining the animal model system, our procedure could be considered an alternative to the complete liquidation of the population in similar cases.

When implementing the rescue procedure we confirmed two facts about PUUV, reported already by other authors [27]: (1) it is not transferred by the placenta and (2) MatAb are transferred via breast milk from mothers and for certain time they protect juveniles from infection. The same appears to also apply to other hantaviruses. For example, in young rats, armed with MatAb and intentionally infected with Seoul orthohantavirus (SEOV; species *Orthohantavirus seoulense*), anti-SEOV-IgM antibody formation peak appears about 2-3 months post infection, whereas in non-MatAb protected animals already about 2-3 weeks post infection [37]. Therefore, we believe the eradication scheme we have conceived and implemented to eradicate PUUV from the bank vole colony can be also effective in the case of other hantaviruses, and perhaps also other viruses infecting other species of animals, provided that the two main characteristics – protection of embryos and protection of juveniles by MatAb – is similarly effective.

## Materials and Methods

### Establishing the colony and the selection experiment

The bank vole colony was reared from wild voles captured in Niepołomice Forest near Kraków (southern Poland) in 2000 and 2001 [48, 49]. In 2004 a multidirectional artificial selection experiment was established, with 4 lines selected for high swim-induced aerobic metabolism, 4 lines selected for herbivorous capability, 4 selected for predatory propensity, and 4 unselected control lines [5, 6]. To preserve genetic polymorphism, in each of the 16 lines the offspring was reared from 16-20 families in each generation (two per year). The selection was effective in each of the three directions, and the unique experimental evolution model system has become a precious resource for the research in several areas, such as behaviour [50, 51], metabolism and thermoregulation [6, 52, 53], molecular genetics [54, 55], exercise physiology [56, 57], neurophysiology [58, 59], ageing [60–62], ecotoxicology [63], stress response [64, 65], holobiont evolution [40, 66] or evolution of dietary niche [67].

Details of the breeding scheme and maintenance conditions are described in our earlier work [6, 50]. Shortly, the colony has been maintained in a “conventional” facility, corresponding to Biosafety Level 1, appropriate for the intended research objectives, which assumed that the animals should not be maintained under germ-free conditions. Depending on the phase of breeding, the colony size ranges from about 1500 to 4000 individuals. The animals are kept in standard plastic mouse cages (1290D or 1264C, Tecniplast, Bugugiatte, Italy) with sawdust bedding, located in six climatic chambers providing fixed light conditions (16L:8D), a constant temperature (20°C) and appropriate ventilation. Food (Labofed H, Morawski, Kcynia, Poland) and water is available *ad libitum*.

### Other voles in the facility

In the same facility, another permanent, but small bank vole colony is maintained, established already in the 1970s, which forms the basis of a separate research programme by our colleagues [68]. The animals are maintained in a partly separate space, which includes animal rooms and laboratory space. However, this space is not separated from the space where the voles from the selection experiment are kept by a sanitary barrier: it is connected to the common corridor by a standard door and uses the same washing and food storage rooms.

In August - November 2009, a separate, temporary colony of voles was established based on animals captured from nine wild populations and used to study variation in physiological performance traits [69, 70]. The animals were kept in a separate room in the same facility as the main colony. The working regime in this colony was separate from that in the main colony (the same person did not work in both colonies on the same day, and separate sets of cages, bottles and other equipment were used), but again the room was not separated by a strict sanitary barrier. Then, two generations of offspring of the wild animals were produced, and finally the entire colony was closed in March 2011. The wild-derived animals were present in the facility from the time when animals of generation 8 of the selection experiment were bred until the time when animals of generation 11 were bred.

### Anaesthesia, euthanasia, and sample collection

For the purpose of the massive and repeated test for the presence of PUUV-specific antibodies (see next section) 5-10µl blood samples were taken from the femoral artery of live animals.

For the purpose of more detailed investigation 40 voles (32 from the infected colony and 8 from the non-infected) were anesthetized with isoflurane (Aerane, Baxter, USA) and blood was collected by cardiac puncture using a heparinized syringe. Voles were then euthanized by administering increasing doses of isoflurane until cardiac arrest, dissected, and the heart and other internal organs were removed (which ensured death). The blood was centrifuged at 4000 rcf for 10 minutes and the plasma was separated. Plasma, blood cells, samples of internal organs, were stored at -80°C (some samples also in RNAlater), and carcasses were stored at -20°C, for further analyses, performed later at the Friedrich-Loeffler-Institut.

### Testing for PUUV presence during the eradication programme

#### Serological tests

Because the eradication plan required repeated testing of the same animals, and getting results as soon as possible after taking the samples, we had to rely on quick, semi-quantitative or qualitative low-cost tests that can be performed on a sample of 5-10μ blood.

The initial tests performed in genP animals, aimed as assessing the age at which all the potential mothers could expect to be seropositive, were performed with a commercial immunochromatographical rapid test based on a purified nucleocapsid protein of PUUV (ReaScan® Ab-Dect Puumala IgG, Reagena, Toivala, Finland; [71]). Blood samples of 10µl were taken from femoral artery blood. The sample was mixed with the dilution buffer with anti-mouse IgG-coated gold particles, and transferred into the test cassette. There the anti-PUUV IgG antibodies will be captured by the membrane-bound PUUV antigen. ReaScan® reader reports the concentration of PUUV-specific antibodies in a numerical, but unitless form. According to the manufacturer instruction, the results were scored as anti-PUUV-negative when <5, and positive when >15.

As the whole eradication plan required performing several thousand tests, for financial reasons the main testing was performed with a low-cost classical IFA (Puumala virus IFA slide, HaartBio Ltd, Helsinki, Finland) [72]), which provides only a qualitative assessment. The IFA tests were used to determine the presence of anti-PUUV antibodies for four purposes: 1) testing the dams from genP after parturition in order to retain only the offspring reared by PUUV-positive ones, 2) testing a group of young voles from genO to assess the age at which MatAb are lost, 3) recurrent, mass-testing of all the individuals from genO (offspring) and genG (grand-offspring) for signs of PUUV infection, 4) testing smaller groups of voles form the neighbouring colony maintained in the same facility, to monitor their infection status.

The tests were performed on 5µl blood sampled from femoral artery, diluted in sterile phosphate-buffered saline (PBS) and stored on ice or in the fridge upon the analysis (performed on the same day). Tests 1, 2 and 4 were performed individually and the blood dilution was 1:10. For the massive tests (test 3), performed repeatedly on all animals, blood samples from the cage-mates were pooled after the sampling in order to increase the analysis throughput and reduce the costs. Typically, the dilutions ranged from 1:4 (when six voles shared the cage) to 1:10 (when only one individual was in the cage). Pilot tests showed that a pool of blood from a single positive and five negative individuals warranted recognizing the sample as positive. When the test was positive for a pool it was repeated for individual voles.

The analyses were performed according to the manufacturer instructions. PUUV-specific antibodies present in PBS-diluted blood or positive control (positive bank vole sera) bind to the PUUV antigen presented on the slides’ wells, after the 30 min of incubation in a humid chamber in 37°C. Not bound antibodies are washed away with PBS and distilled water. Fluorescein isothiocyanate (FITC) conjugated to polyclonal rabbit anti-mouse IgG (product F0261, Dako, Glostrup, Denmark) was added to all the wells to enable the antigen-antibody bond detection under the light microscope with fluorescence excitation (Eclipse 80i, Nikon Instruments Inc., Melville NY, USA). The wells were rated independently just after completing the slide processing.

Several preliminary trials with the IFA test were performed to establish a grading scale based on the brightness of the fluorescence, and the scale was learned by all the persons involved in the testing. Initially, the scale had 6 grades (from 0 = “certainly negative” to 5 = “strongly positive”), but then it was reduced to three-levels: 0 = negative, 1 = doubtful, and 2 = positive. During the main work, each slide was judged independently by two observers. If both observers scored the sample as either positive or negative, the result was taken as confirmed. If their scores differed, or both scored the result as doubtful, the test was repeated on the next day on a new blood sample. As the positive and negative control for the IFA tests we used mixed samples of blood collected at the time of euthanasia (and stored at -80°C) from the animals of genP, which were tested earlier with the ReaScan® test.

#### Viral RNA detection

To detect infected individuals from genO of the eradication programme, samples of saliva from alive individuals and of dust from their cages were taken for the detection of viral RNA. To confirm the infection in the genO animals that appeared anti-PUUV-positive in IFA tests, and to confirm the absence of PUUV at the final stage of the eradication programme, the animals were euthanized (cervical dislocation or CO_2_ inhalation) and dissected, and lung samples were taken and placed into RNAlater. Before the RNA extraction, the tissue was homogenized using motor homogenizer with diethyl pyrocarbonate (DEPC)-treated metal tips (2 x 15 sec). Additionally, samples of dust from various places of the animal facility were also analysed to confirm the efficiency of the facility disinfection.

RNA was extracted using RNAzol® (Molecular Research Center, Cincinnati, OH, USA). To detect hantavirus RNA, a quantitative reverse transcription-polymerase chain reaction (RT-qPCR) was performed according the QuantiTect® Probe RT-PCR Kit protocol (Qiagen, Venlo, Netherlands), using the primers F (5’-gtgcaccagatcgrtgtcc-3’) and R (5’-yarctctgccatccctgca-3’) and the TaqMan® probe (5’-ccaacatgyatttatg-3’) (Syngen Biotech, Wrocław, Poland) (a protocol of [23], modified according to personal advise from Liina Voutilainen). The amplification was performed with 7500 Fast Real-Time PCR System (Applied Biosystems Inc, Foster City, CA, USA). The amplification programme consisted of one cycle of an initial incubation at 50°C for 30min and denaturation at 95°C for 15min, followed by 45 cycles of annealing at 94°C for 15s, and extension at 59°C for 60s [23]. Each set of analyses was run with a negative (all reagents without RNA sample) and a positive control (pooled RNA extracted from lung samples of several voles with known anti-PUUV IgG level, identified at the beginning of the eradication procedure). The result was considered positive if the detectable signal from the probe was achieved, and it was expressed as the number of a cycle when threshold was exceeded (Cт).

### Identification of the putative PUUV origin

#### Serological investigations

Chest cavity lavage (CCL) samples were analysed using the same commercial rapid test and an in-house IgG enzyme-linked immunosorbent assay (ELISA) as previously described [71, 73]. As a positive control the monoclonal antibody (mAb) 5E11 was used [74]. CCL of a bank vole tested negative before by serological and molecular methods and originating from a region where PUUV is absent, was used as negative control.

#### Molecular investigations

For identification of the origin of PUUV incursion, RNA was isolated from a lentil-sized piece of lung tissue from 22 bank voles using an in-house protocol and QIAzol® Lysis Reagent (QIAGEN, Hilden, Germany) [75]. Conventional RT-PCR assays of the S segment, M segment and L segment were applied as described before [73, 76, 77] using the SuperScript™ III One-Step RT-PCR with Platinum Taq-Kit (Invitrogen, Darmstadt, Germany). RT-PCR products of the expected size were sequenced by the dideoxy-chain termination method with the BigDye Terminator v1.1 Kit (Applied Biosystems, Darmstadt, Germany). Sequence alignments were established with the software BioEdit v7.2.5 as well as pairwise sequence comparison [78]. The most suitable substitution model was determined by jModelTest v2.1.8 [79]. Phylogentic trees were reconstructed via MrBayes v3.2.6 [80, 81].

### Sanitary precautions

All works connected with standard animal care were performed in controlled conditions, minimising virus transmission. Manipulations requiring opening the cage (i.e., changing cages, providing food, taking samples) were performed using cage changing station with laminar airflow and HEPA filters (model CS5, Tecniplast, Bugugiatte, Italy). All tools, equipment, working surfaces, chambers, corridor and facility rooms (storage, cleaning etc.) were continually disinfected with antiviral agent (Virkon; Naturan, Warsaw, Poland). The entire ventilation system in emptied rooms were disinfected soon after moving juveniles into the IVC-system.

## Ethical statement

Animal welfare was monitored daily throughout the experiment. The procedures on animals performed as a part of the maintenance of the experimental colony were approved by the 1st Local Institutional Animal Care and Use Committee (decision 68/2012), in accordance with the EU directive 2010/63/EU. According the regulations effective in 2013, the additional procedure of blood sampling performed exclusively for the purpose of the PUUV detection, rather than as a part of an experiment performed for research purposes, did not require a separate ethical permit.

## Acknowledgements

We are grateful to: Students and technicians who worked with the voles and helped with testing (Agata Rudolf, Geoffrey Dheyongera, Maciej Działo, and Michał Kolasa); Director of the Institute of Environmental Sciences and the Dean of the Faculty of Biology of the Jagiellonian University for granting special funds for equipment, and the University administration for enabling fast-track purchases; Polish National Science Centre for the agreement to redirect funds from grants to cover the costs of the eradication rescuing the colony; The Veterinary and Sanitary Inspection Officers for granting us permission to attempt to rescue the colony; Florian Binder, Kathrin Jeske, and Dörte Kaufmann for help in identification of the PUUV origin of the outbreak in the colony; Olli Vapalahti for IFA slides protocol, and Liina Voutilainen for help with protocols of RNA analyses; The building management staff for their cooperation and servicing all the systems needed to run the facility during the infection outbreak; Businesses: Anima and Tecniplast for offering a discount on the equipment and for arranging fast-track delivery.

## Supplementary Information

**S1 Fig. Phylogenetic trees of partial Puumala virus (PUUV) sequences from the laboratory (lab.) colony and reference sequences from Poland and other representative strains.** The sequences are from A) the S segment with 711nt, B) M segment with 618nt, and C) L segment with 411nt in length. The consensus sequences and alignments were constructed with BioEdit v7.2.5 [82] and the best substitution model determined with JModelTest v2.1.8 [79]. The phylogenetic trees were calculated with the aid of MrBayes v3.2.6 with up to 4 x 10^6^ generations, a burn in of 25% and two-parameter substitution models with gamma distribution and invariant sites [80]. PUUV lineages: ALAD Alpe-Adrian, CE Central European, DAN Danish, FIN Finnish, LAT Latvian, N-SCA North-Scandinavian, RUS Russian, S-SCA South-Scandinavian. Outgroups: MUJV Muju virus, TULV Tula virus. {References are in the supplement].

**S1 Table. Serological and RT-PCR analyses of 22 bank voles of the generation 14 of the bank vole colony.**

**S2 Table. Pairwise nucleotide (nt) and amino acid (aa) sequence similarities of partial S (a), M (b) and L (c) S segment sequences of Puumala virus (PUUV) detected in the colony, reference sequences from Poland, and other representative strains of PUUV clades.**

## Data Availability Statement

Data that support the findings of this study are available in the Materials and Methods, Results, and/or Supplemental Material of this article.

## Author Contributions

J. Hajduk proposed the initial idea of the eradication plan; P. Koteja, with the help of E.T. Sadowska, J. Wyszkowska, E. Jaromin, J. Hajduk, I. Onik, B. Bober-Sowa and R.G. Ulrich designed the detailed plan of the eradication; E. Sadowska, K. Baliga-Klimczyk and B. Bober-Sowa supervised the animal colony; B. Bober-Sowa, K. Baliga-Klimczyk, J. Hajduk, I. Onik, K. Lach, J. Wyszkowska, E. Jaromin and E.T. Sadowska performed the animal maintenance work and most of the tests; M. Konczal refined the protocol of RT-qPCR and performed most of these tests; S. Drewes and R.G. Ulrich performed the serological and molecular analyses for the purpose of establishing the PUUV origin and phylogeny; J. Wyszkowska, E. Jaromin and S. Drewes drafted the manuscript; P. Koteja and S. Drewes, with the help of R.G. Ulrich, wrote the final manuscript; All authors commented on the final version of the text.

## References

1. Traub E. The epidemiology of lymphocytic choriomeningitis in white mice. J Exp Med. 1936;64(2):183–200.

2. Iwai H, Itoh T, Shumiya S. Persistence of Sendai virus in a mouse breeder colony and possibility to re-establish the virus free colonies. Jikken Dobutsu. 1977;26(3):205–12.

3. Hotchin J, Sikora E, Kinch W, Hinman A, Woodall J. Lymphocytic choriomeningitis in a hamster colony causes infection of hospital personnel. Science. 1974;185(4157):1173-4.

4. Knust B, Brown S, de St. Maurice A, Whitmer S, Koske SE, Ervin E, et al. Seoul Virus Infection and Spread in United States Home-Based Ratteries: Rat and Human Testing Results From a Multistate Outbreak Investigation. J Infect Dis. 2020;222(8):1311–9.

5. Sadowska ET, Baliga-Klimczyk K, Chrząścik KM, Koteja P. Laboratory model of adaptive radiation: a selection experiment in the bank vole. Physiol Biochem Zool. 2008;81(5):627–40.

6. Sadowska ET, Stawski C, Rudolf A, Dheyongera G, Chrzascik KM, Baliga-Klimczyk K, et al. Evolution of basal metabolic rate in bank voles from a multidirectional selection experiment. Proc Biol Sci. 2015;282(1806):20150025.

7. Bujalska G, Hansson L. Bank vole biology: recent advances in the population biology of a model species. Pol J Ecol. 2000;48(Suppl.).

8. Boratyński Z, Szyrmer M, Koteja P. The metabolic performance predicts home range size of bank voles: a support for the behavioral-bioenergetics theory. Oecologia. 2020;193(3):547–56.

9. Brila I, Lavrinienko A, Tukalenko E, Ecke F, Rodushkin I, Kallio ER, et al. Low-level environmental metal pollution is associated with altered gut microbiota of a wild rodent, the bank vole (Myodes glareolus). Sci Total Environ. 2021;790:148224.

10. McManus A, Holland CV, Henttonen H, Stuart P. The Invasive Bank Vole (Myodes glareolus): A Model System for Studying Parasites and Ecoimmunology during a Biological Invasion. Animals (Basel). 2021;11(9).

11. Röhrs S, Begeman L, Straub BK, Boadella M, Hanke D, Wernike K, et al. The bank vole (*Clethrionomys glareolus*) - small animal model for hepacivirus infection. Viruses. 2021;13(12):2421.

12. Kauer L, Imholt C, Jacob J, Berens C, Kühn R. Seasonal shifts and land-use impact: unveiling the gut microbiomes of bank voles (*Myodes glareolus*) and common voles (*Microtus arvalis*). FEMS Microbiol Ecol. 2024;100(12).

13. Paul L, Kirsch P, Thomzig A, Thone-Reineke C, Beekes M. Practical approaches for refinement and reduction of animal experiments with bank voles in prion research. Berliner Und Munchener Tierarztliche Wochenschrift. 2018;131(9-10):359–67.

14. Jeske K, Weber S, Pfaff F, Imholt C, Jacob J, Beer M, et al. Molecular Detection and Characterization of the First Cowpox Virus Isolate Derived from a Bank Vole. Viruses. 2019;11(11).

15. Michelitsch A, Tews BA, Klaus C, Bestehorn-Willmann M, Dobler G, Beer M, et al. In Vivo Characterization of Tick-Borne Encephalitis Virus in Bank Voles (*Myodes glareolus*). Viruses. 2019;11(11).

16. Heyman P, Ceianu CS, Christova I, Tordo N, Beersma M, João Alves M, et al. A five-year perspective on the situation of haemorrhagic fever with renal syndrome and status of the hantavirus reservoirs in Europe, 2005-2010. Euro Surveill. 2011;16(36):pii=19961.

17. Fischer S, Mayer-Scholl A, Imholt C, Spierling NG, Heuser E, Schmidt S, et al. Leptospira Genomospecies and Sequence Type Prevalence in Small Mammal Populations in Germany. Vector Borne Zoonotic Dis. 2018;18(4):188–99.

18. Nainys J, Timinskas A, Schneider J, Ulrich RG, Gedvilaite A. Identification of two novel members of the tentative genus Wukipolyomavirus in wild rodents. PLoS One. 2015;10(10):e0140916.

19. Drexler JF, Corman VM, Müller MA, Lukashev AN, Gmyl A, Coutard B, et al. Evidence for novel hepaciviruses in rodents. PLoS Pathog. 2013;9(6):e1003438.

20. Heyman P, Cochez C, Ducoffre G, Mailles A, Zeller H, Abu Sin M, et al. Haemorrhagic Fever with Renal Syndrome: an analysis of the outbreaks in Belgium, France, Germany, the Netherlands and Luxembourg in 2005. Euro Surveill. 2007;12(5):E15–6.

21. Avšič-Županc T, Saksida A, Korva M. Hantavirus infections. Clin Microbiol Infect. 2019;21:e6–e16.

22. Vapalahti O, Mustonen J, Lundkvist Å, Henttonen H, Plyusnin A, Vaheri A. Hantavirus infections in Europe. Lancet Infect Dis. 2003;3(10):653–61.

23. Hardestam J, Karlsson M, Falk KI, Olsson G, Klingström J, Lundkvist Å. Puumala hantavirus excretion kinetics in bank voles (*Myodes glareolus*). Emerg Infect Dis. 2008;14(8):1209–15.

24. Kallio ER, Klingström J, Gustafsson E, Manni T, Vaheri A, Henttonen H, et al. Prolonged survival of Puumala hantavirus outside the host: evidence for indirect transmission via the environment. J Gen Virol. 2006;87(Pt 8):2127–34.

25. Yanagihara R, Amyx HL, Gajdusek DC. Experimental infection with Puumala virus, the etiologic agent of nephropathia epidemica, in bank voles (*Clethrionomys glareolus*). J Virol. 1985;55(1):34–8.

26. Gavrilovskaya IN, Apekina NS, Bernshtein AD, Varvara TD, Okulova NM, Myasnikov YA, et al. Pathogenesis of hemorrhagic fever with renal syndrome virus infection and mode of horizontal transmission of hantavirus in bank voles. In: Calisher CH, editor. Hemorrhagic Fever with Renal Syndrome, Tick- and Mosquito-Borne Viruses Archives of Virology Supplementum I. Wien: Springer-Verlag; 1990. p. 57–62.

27. Kallio ER, Poikonen A, Vaheri A, Vapalahti O, Henttonen H, Koskela E, et al. Maternal antibodies postpone hantavirus infection and enhance individual breeding success. Proc R Soc B. 2006;273(1602):2771-6.

28. Botten J, Mirowsky K, Kusewitt D, Bharadwaj M, Yee J, Ricci R, et al. Experimental infection model for Sin Nombre hantavirus in the deer mouse (*Peromyscus maniculatus*). Proc Natl Acad Sci U S A. 2000;97(19):10578–83.

29. Krüger DH, Schönrich G, Klempa B. Human pathogenic hantaviruses and prevention of infection. Hum Vaccin. 2011;7(6):685–93.

30. Reil D, Rosenfeld UM, Imholt C, Schmidt S, Ulrich RG, Eccard JA, et al. Puumala hantavirus infections in bank vole populations: host and virus dynamics in Central Europe. BMC Ecol. 2017;17(1):9.

31. Reil D, Imholt C, Eccard JA, Jacob J. Beech fructification and bank vole population dynamics - combined analyses of promoters of human Puumala Virus infections in Germany. PLoS One. 2015;10(7):e0134124.

32. Baumann A, Dudek D, Sadkowska-Todys M. [The role of natural environment in spreading of hantavirus--model of the correlation between host, pathogen and human infections]. Przegl Epidemiol. 2007;61(4):647–55.

33. Tersago K, Verhagen R, Servais A, Heyman P, Ducoffre G, Leirs H. Hantavirus disease (nephropathia epidemica) in Belgium: effects of tree seed production and climate. Epidemiol Infect. 2009;137(2):250–6.

34. Knust B, Stroher U, Edison L, Albarino CG, Lovejoy J, Armeanu E, et al. Lymphocytic choriomeningitis virus in employees and mice at multipremises feeder-rodent operation, United States, 2012. Emerg Infect Dis. 2014;20(2):240–7.

35. Voutilainen L, Sironen T, Tonteri E, Bäck AT, Razzauti M, Karlsson M, et al. Life-long shedding of Puumala hantavirus in wild bank voles (*Myodes glareolus*). J Gen Virol. 2015;96(Pt 6):1238–47.

36. Kim H, Bang J, Baek SH, Park JH. Eliminating murine norovirus, Helicobacter hepaticus, and intestinal protozoa by embryo transfer for an entire mouse barrier facility. Exp Anim. 2022;71(1):28–35.

37. Dohmae K, Nishimune Y. Protection against hantavirus infection by dam’s immunity transferred vertically to neonates. Arch Virol. 1995;140(1):165–72.

38. Artwohl JE, Purcell JE, Fortman JD. The use of cross-foster rederivation to eliminate murine norovirus, Helicobacter spp., and murine hepatitis virus from a mouse colony. J Am Assoc Lab Anim Sci. 2008;47(6):19–24.

39. Buxbaum LU, DeRitis PC, Chu N, Conti PA. Eliminating murine norovirus by cross-fostering. J Am Assoc Lab Anim Sci. 2011;50(4):495–9.

40. Lipowska MM, Sadowska ET, Kohl KD, Koteja P. Experimental Evolution of a Mammalian Holobiont? Genetic and Maternal Effects on the Cecal Microbiome in Bank Voles Selectively Bred for Herbivorous Capability. Ecol Evol Physiol. 2024;97(5):274–91.

41. Van Epps HL, Terajima M, Mustonen J, Arstila TP, Corey EA, Vaheri A, et al. Long-lived memory T lymphocyte responses after hantavirus infection. J Exp Med. 2002;196(5):579–88.

42. Weber de Melo V, Sheikh Ali H, Freise J, Kühnert D, Essbauer S, Mertens M, et al. Spatiotemporal dynamics of Puumala hantavirus associated with its rodent host, *Myodes glareolus*. Evol Appl. 2015;8(6):545–59.

43. Binder F, Ryll R, Drewes S, Jagdmann S, Reil D, Hiltbrunner M, et al. Spatial and Temporal Evolutionary Patterns in Puumala Orthohantavirus (PUUV) S Segment. Pathogens. 2020;9(7):548.

44. Filipi K, Marková S, Searle JB, Kotlík P. Mitogenomic phylogenetics of the bank vole *Clethrionomys glareolus*, a model system for studying end-glacial colonization of Europe. Mol Phylogenet Evol. 2015;82 Pt A:245–57.

45. Drewes S, Ali HS, Saxenhofer M, Rosenfeld UM, Binder F, Cuypers F, et al. Host-associated absence of human Puumala Virus infections in northern and eastern Germany. Emerg Infect Dis. 2017;23(1):83–6.

46. Ali HS, Drewes S, Sadowska ET, Mikowska M, Groschup MH, Heckel G, et al. First molecular evidence for Puumala hantavirus in Poland. Viruses. 2014;6(1):340–53.

47. Rosenfeld UM, Drewes S, Ali HS, Sadowska ET, Mikowska M, Heckel G, et al. A highly divergent Puumala virus lineage in southern Poland. Arch Virol. 2017;162(5):1177–85.

48. Labocha MK, Sadowska ET, Baliga K, Semer AK, Koteja P. Individual variation and repeatability of basal metabolism in the bank vole, Clethrionomys glareolus. Proc Biol Sci. 2004;271(1537):367-72.

49. Sadowska ET, Baliga-Klimczyk K, Labocha MK, Koteja P. Genetic Correlations in a Wild Rodent: Grass-Eaters and Fast-Growers Evolve High Basal Metabolic Rates. Evolution. 2009;63(6):1530–9.

50. Chrząścik KM, Sadowska ET, Rudolf A, Koteja P. Learning ability in bank voles selected for high aerobic metabolism, predatory behaviour and herbivorous capability. Physiol Behav. 2014;135:143–51.

51. Maiti U, Sadowska ET, ChrzAscik KM, Koteja P. Experimental evolution of personality traits: open-field exploration in bank voles from a multidirectional selection experiment. Curr Zool. 2019;65(4):375–84.

52. Sadowska ET, Krol E, Chrzascik KM, Rudolf AM, Speakman JR, Koteja P. Limits to sustained energy intake. XXIII. Does heat dissipation capacity limit the energy budget of lactating bank voles? J Exp Biol. 2016;219:805–15.

53. Stawski C, Koteja P, Sadowska ET, Jefimow M, Wojciechowski MS. Selection for high activity-related aerobic metabolism does not alter the capacity of non-shivering thermogenesis in bank voles. Comp Biochem Physiol A Mol Integr Physiol. 2015;180:51–6.

54. Konczal M, Babik W, Radwan J, Sadowska ET, Koteja P. Initial Molecular-Level Response to Artificial Selection for Increased Aerobic Metabolism Occurs Primarily through Changes in Gene Expression. Mol Biol Evol. 2015;32(6):1461–73.

55. Konczal M, Koteja P, Orlowska-Feuer P, Radwan J, Sadowska ET, Babik W. Genomic Response to Selection for Predatory Behavior in a Mammalian Model of Adaptive Radiation. Mol Biol Evol. 2016;33(9):2429–40.

56. Jaromin E, Wyszkowska J, Labecka AM, Sadowska ET, Koteja P. Hindlimb muscle fibre size and glycogen stores in bank voles with increased aerobic exercise metabolism. J Exp Biol. 2016;219(Pt 4):470–3.

57. Lipowska MM, Dheyongera G, Sadowska ET, Koteja P. Experimental evolution of aerobic exercise performance and hematological traits in bank voles. Comparative Biochemistry and Physiology a-Molecular & Integrative Physiology. 2019;234:1–9.

58. Jaromin E, Sadowska ET, Koteja P. The effect of monoamines reuptake inhibitors on aerobic exercise performance in bank voles from a selection experiment. Curr Zool. 2019;65(4):409–19.

59. Jaromin E, Sadowska ET, Koteja P. Is Experimental Evolution of an Increased Aerobic Exercise Performance in Bank Voles Mediated by Endocannabinoid Signaling Pathway? Front Physiol. 2019;10:640.

60. Rudolf AM, Dańko MJ, Sadowska ET, Dheyongera G, Koteja P. Age-related changes of physiological performance and survivorship of bank voles selected for high aerobic capacity. Exp Gerontol. 2017;98:70–9.

61. Grosiak M, Koteja P, Bauchinger U, Sadowska ET. Age-Related Changes in the Thermoregulatory Properties in Bank Voles From a Selection Experiment. Front Physiol. 2020;11:576304.

62. Grosiak M, Koteja P, Hambly C, Speakman JR, Sadowska ET. Limits to sustained energy intake. XXXIV. Can the heat dissipation limit (HDL) theory explain reproductive aging? J Exp Biol. 2024;227(4).

63. Dheyongera G, Grzebyk K, Rudolf AM, Sadowska ET, Koteja P. The effect of chlorpyrifos on thermogenic capacity of bank voles selected for increased aerobic exercise metabolism. Chemosphere. 2016;149:383–90.

64. Lipowska MM, Sadowska ET, Bauchinger U, Goymann W, Bober-Sowa B, Koteja P. Does selection for behavioral and physiological performance traits alter glucocorticoid responsiveness in bank voles? J Exp Biol. 2020;223(Pt 15).

65. Lipowska MM, Sadowska ET, Palme R, Koteja P. Evolution of an increased performance under acute challenge does not exacerbate vulnerability to chronic stress. Sci Rep. 2022;12(1):2126.

66. Hanhimäki E, Watts PC, Koskela E, Koteja P, Mappes T, Hämäläinen AM. Evolved high aerobic capacity has context-specific effects on gut microbiota. Frontiers in Ecology and Evolution. 2022;10.

67. Hämäläinen A, Kiljunen M, Koskela E, Koteja P, Mappes T, Rajala M, et al. Artificial selection for predatory behaviour results in dietary niche differentiation in an omnivorous mammal. Proc Biol Sci. 2022;289(1970):20212510.

68. Kapusta J, Kruczek M, Pochroń E, Olejniczak P. Maintenance of seasonal differences in reproductive characteristics of bank voles after 30 years captive breeding. Scandinavian Journal of Laboratory Animal Science. 2012;39(1):91–100.

69. Mikowska M, Gaura A, Sadowska E, Koteja P, Świergosz-Kowalewska R. Genetic variation in bank vole populations in natural and metal-contaminated areas. Arch Environ Contam Toxicol. 2014;67(4):535–46.

70. Sadowska ET, Mikowska M, Świergosz-Kowalewska R, Koteja P. Physiological performance in the small rodent Myodes glareolus from isolated and heavily polluted populations. Programme and Abstract Book. Prague, Czech Republic: Annual Main Meeting of the Society for Experimental Biology; 2010. p. 209–10.

71. Sirola H, Kallio ER, Koistinen V, Kuronen I, Lundkvist Å, Vaheri A, et al. Rapid field test for detection of hantavirus antibodies in rodents. Epidemiol Infect. 2004;132(3):549–53.

72. Hedman K, Vaheri A, Brummer-Korvenkontio M. Rapid diagnosis of hantavirus disease with an IgG-avidity assay. Lancet. 1991;338(8779):1353-6.

73. Mertens M, Kindler E, Emmerich P, Esser J, Wagner-Wiening C, Wölfel R, et al. Phylogenetic analysis of Puumala virus subtype Bavaria, characterization and diagnostic use of its recombinant nucleocapsid protein. Virus Genes. 2011;43(2):177–91.

74. Žvirblienė A, Samonskyte L, Gedvilaitė A, Voronkova T, Ulrich R, Sasnauskas K. Generation of monoclonal antibodies of desired specificity using chimeric polyomavirus-derived virus-like particles. J Immunol Methods. 2006;311(1-2):57–70.

75. Schmidt S, Saxenhofer M, Drewes S, Schlegel M, Wanka KM, Frank R, et al. High genetic structuring of Tula hantavirus. Arch Virol. 2016;161(5):1135–49.

76. Essbauer S, Schmidt J, Conraths FJ, Friedrich R, Koch J, Hautmann W, et al. A new Puumala hantavirus subtype in rodents associated with an outbreak of Nephropathia epidemica in South-East Germany in 2004. Epidemiol Infect. 2006;134(6):1333–44.

77. Klempa B, Fichet-Calvet E, Lecompte E, Auste B, Aniskin V, Meisel H, et al. Hantavirus in African wood mouse, Guinea. Emerg Infect Dis. 2006;12(5):838–40.

78. Hall T. BioEdit: an important software for molecular biology. GERF Bull Biosci. 2011;2:60–1.

79. Darriba D, Taboada GL, Doallo R, Posada D. jModelTest 2: more models, new heuristics and parallel computing. Nat Methods. 2012;9(8):772.

80. Ronquist F, Teslenko M, van der Mark P, Ayres DL, Darling A, Höhna S, et al. MrBayes 3.2: efficient Bayesian phylogenetic inference and model choice across a large model space. Syst Biol. 2012;61(3):539–42.

81. Price MN, Dehal PS, Arkin AP. FastTree 2 - approximately maximum-likelihood trees for large alignments. PLoS One. 2010;5(3):e9490.

82. Hall TA. BioEdit: a user-friendly biological sequence alignment editor and analysis program for Windows 95/98/NT. Nucl Acids Symp Ser. 1999;41(41):95–8.

