## Supplementary Figure 1 for "Maternal antibody-mediated elimination of a Puumala hantavirus outbreak in a bank vole colony"

**Supplementary S1 Figure**

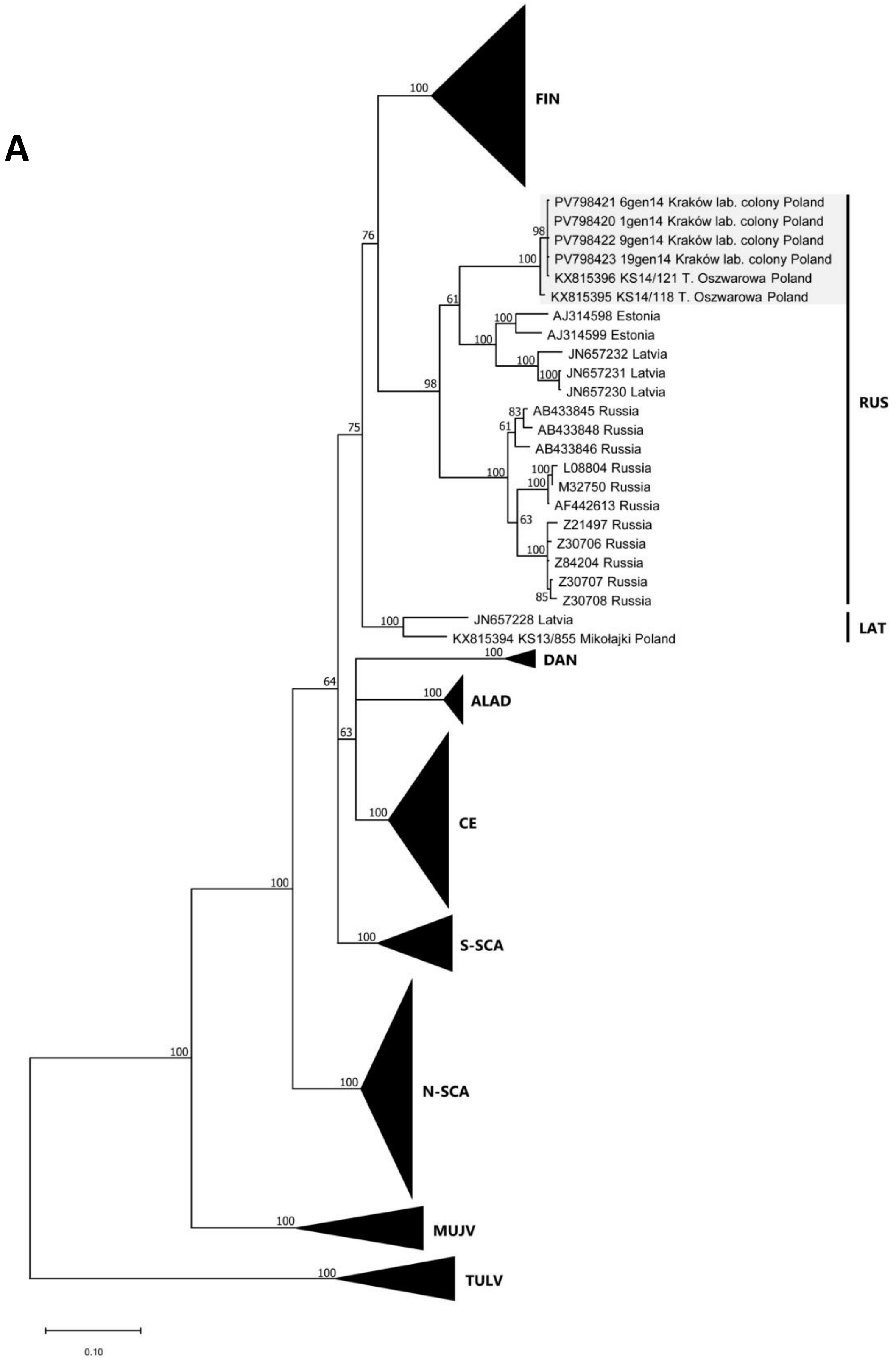

B

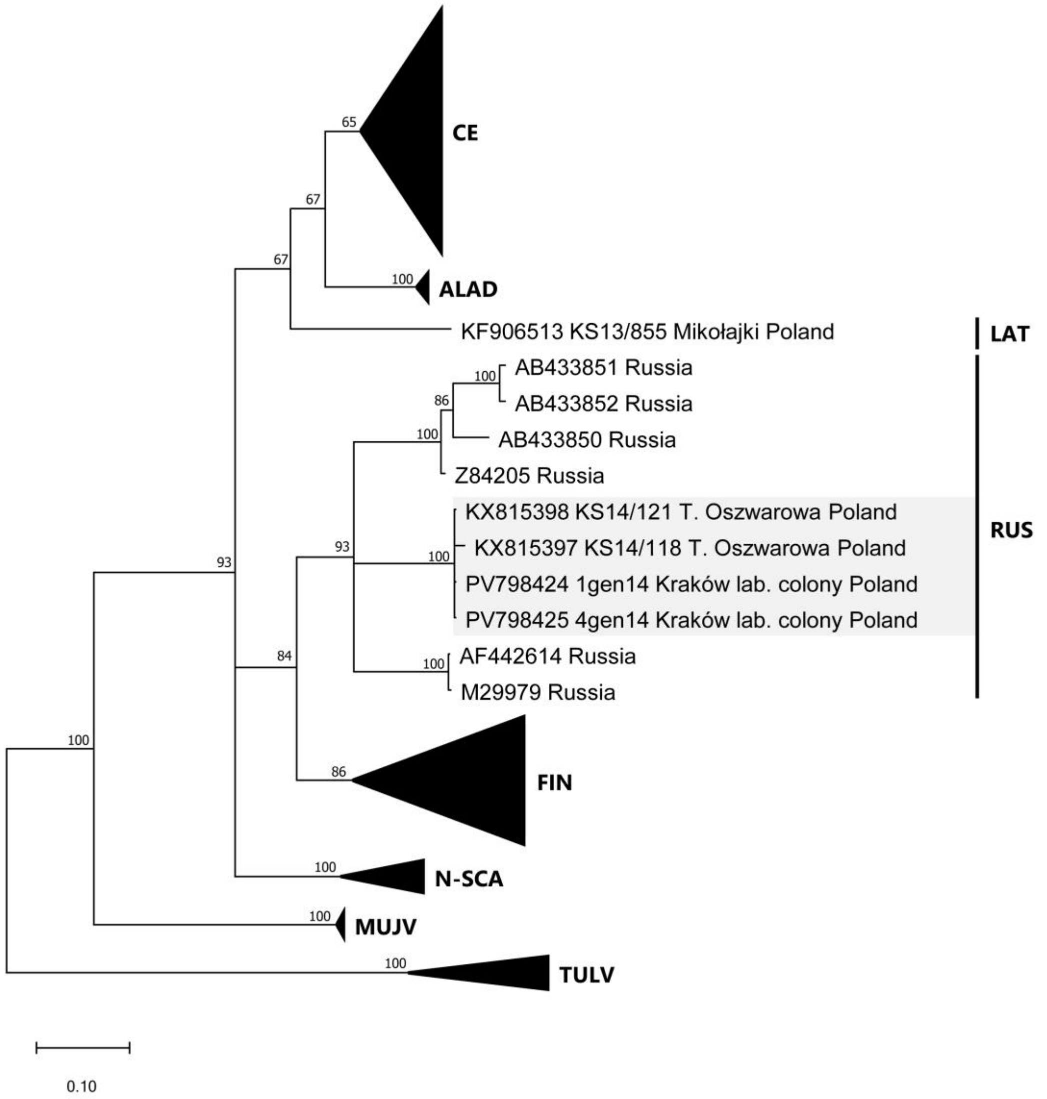
CE
ALAD
KF906513 KS13/855 Mikołajki Poland
LAT
AB433851 Russia
AB433852 Russia
AB433850 Russia
Z84205 Russia
KX815398 KS14/121 T. Oszwarowa Poland
KX815397 KS14/118 T. Oszwarowa Poland
PV798424 1gen14 Kraków lab. colony Poland
PV798425 4gen14 Kraków lab. colony Poland
AF442614 Russia
M29979 Russia
RUS
FIN
N-SCA
MUJV
TULV
0.10

C

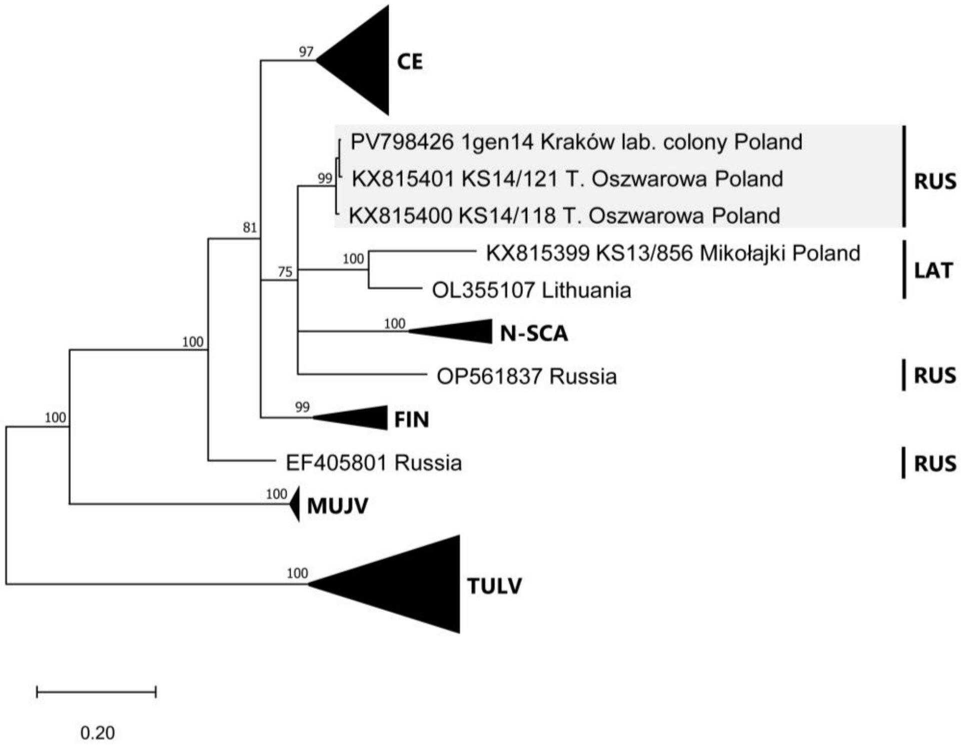


**S1 Fig. Phylogenetic trees of partial Puumala virus (PUUV) sequences from the laboratory (lab.) colony and reference sequences from Poland and other representative strains.** The sequences are from A) the S segment with 711nt, B) M segment with 618nt, and C) L segment with 411nt in length. The consensus sequences and alignments were constructed with BioEdit v7.2.5 [1] and the best substitution model determined with JModelTest v2.1.8 [2]. The phylogenetic trees were calculated with the aid of MrBayes v3.2.6 with up to 4 x $10^6$ generations, a burn in of 25% and two-parameter substitution models with gamma distribution and invariant sites [3]. PUUV lineages: ALAD Alpe-Adrian, CE Central European, DAN Danish, FIN Finnish, LAT Latvian, N-SCA North-Scandinavian, RUS Russian, S-SCA South-Scandinavian. Outgroups: MUJV Muju virus, TULV Tula virus.

### References


1. T. A. Hall "BioEdit: a user-friendly biological sequence alignment editor and analysis program for Windows 95/98/NT," *Nucleic acids symposium series* 41 (1999): 95-98

2. D. Darriba, G. L. Taboada, R. Doallo, and D. Posada "jModelTest 2: more models, new heuristics and parallel computing," *Nature methods* 9 (2012): 772

3. F. Ronquist, M. Teslenko, P. van der Mark, D. L. Ayres, A. Darling, S. Höhna, et al., "MrBayes 3.2: efficient Bayesian phylogenetic inference and model choice across a large model space," *Systematic biology* 61 (2012): 539-542
