## Supplementary Table 1 for "Maternal antibody-mediated elimination of a Puumala hantavirus outbreak in a bank vole colony"

**S1 Table.** Serological and RT-PCR analyses of 17 bank voles of the generation 14 (Z) of the bank vole colony and five bank voles from a separate colony kept in different room (MRT).

| IR | ID | PUUV IgG ELISA | PUUV S-RT-PCR | PUUV Rapid test Reascan |
|---|---|---|---|---|
| 1 | Z56527 | pos | pos | pos |
| 2 | Z56091 | pos | pos | pos |
| 3 | MRT | neg | neg | neg |
| 4 | Z56058 | pos | pos | pos |
| 5 | Z55488 | pos | pos | pos |
| 6 | Z60057 | pos | pos | pos |
| 7 | MRT | neg | neg | neg |
| 8 | Z56771 | pos | pos | pos |
| 9 | Z60031 | pos | pos | pos |
| 10 | Z56373 | pos | pos | pos |
| 11 | MRT | neg | neg | neg |
| 12 | Z60222 | pos | pos | pos |
| 13 | Z55491 | pos | pos | pos |
| 14 | Z53904 | pos | pos | pos |
| 15 | Z54860 | pos | pos | pos |
| 16 | MRT | neg | neg | neg |
| 17 | Z55666 | pos | pos | pos |
| 18 | Z55547 | pos | pos | pos |
| 19 | Z53707 | pos | pos | pos |
| 20 | Z56263 | pos | pos | pos |
| 21 | MRT | neg | neg | neg |
| 22 | Z56466 | pos | pos | equ |
| total | | 17/22 | 17/22 | 16/22 |

IR record number, ID identification number, PUUV Puumala virus, IgG ELISA immunoglobulin G enzyme-linked immunosorbent assay, S-RT-PCR reverse transcription-polymerase chain reaction targeting the small (S) segment, pos positive, neg negative, equ equivocal.
