## Supplementary Table 2 for "Maternal antibody-mediated elimination of a Puumala hantavirus outbreak in a bank vole colony"

**S2 Table.** Pairwise nucleotide (nt) and amino acid (aa) sequence similarities of partial S (a: 711 nt length), M (b: 618 nt length) and L (c: 411 nt length) segment sequences of Puumala virus (PUUV) detected in generation 14 (gen14) of the colony, reference sequences from Poland, and other representative strains of PUUV clades.

| (a) Reference sequences of the S segment | PV798420 1gen14 Kraków Poland (RUS) | | PV798421 6gen14 Kraków Poland (RUS) | | PV798422 9gen14 Kraków Poland (RUS) | | PV798423 19gen14 Kraków Poland (RUS) | |
|---|---|---|---|---|---|---|---|---|
| | nt | aa | nt | aa | nt | aa | nt | aa |
| PV798420 1gen14 Kraków lab. colony Poland (RUS) | - | - | **0.998** | 1 | **0.998** | 1 | **0.998** | 0.995 |
| PV798421 6gen14 Kraków lab. colony Poland (RUS) | **0.998** | 1 | - | - | 0.997 | 1 | 0.997 | 0.995 |
| PV798422 9gen14 Kraków lab. colony Poland (RUS) | **0.998** | 1 | 0.997 | 1 | - | - | 0.997 | 0.995 |
| PV798423 19gen14 Kraków lab. colony Poland (RUS) | **0.998** | 0.995 | 0.997 | 0.995 | 0.997 | 0.995 | - | - |
| KX815395 KS14/118 Teleśnica Oszwarowa Poland (RUS) | 0.987 | 0.991 | 0.985 | 0.991 | 0.985 | 0.991 | 0.985 | 0.987 |
| KX815396 KS14/121 Teleśnica Oszwarowa Poland (RUS) | **0.998** | 0.995 | 0.997 | 0.995 | 0.997 | 0.995 | 0.997 | 0.991 |
| KX815394 KS13/855 Mikołajki Poland (LAT) | 0.835 | 0.978 | 0.834 | 0.978 | 0.834 | 0.978 | 0.835 | 0.974 |
| JN657228 Latvia (LAT) | 0.838 | 0.978 | 0.839 | 0.978 | 0.836 | 0.978 | 0.838 | 0.974 |
| JN657230 Latvia (RUS), JN657231, JN657232 | 0.872-0.876 | 0.974 | 0.870-0.874 | 0.974 | 0.870-0.874 | 0.974 | 0.870-0.874 | 0.970 |
| AJ314598 Estonia (RUS), AJ314599 | 0.856-0.864 | 0.987-0.991 | 0.855-0.863 | 0.987-0.991 | 0.855-0.863 | 0.987-0.991 | 0.856-0.864 | 0.983-0.987 |
| AF442613 Russia (RUS), M32750, L08804, Z84204, AB433846, AB433848, AB433845, Z30708, Z30706, Z30707, Z21497 | 0.846-0.863 | 0.970-0.978 | 0.845-0.862 | 0.970-0.978 | 0.845-0.862 | 0.970-0.978 | 0.846-0.863 | 0.966-0.974 |
| Z30702 Finland (FIN), Z30704, Z30705, GU808825, GU808824, AJ314597, Z46942, Z69985, JQ319171, JQ319163, JQ319166, JQ319167, JQ319168, JQ319161, JQ319164, JQ319170, JQ319162, JN831950, HE801633 | 0.815-0.832 | 0.928-0.966 | 0.817-0.831 | 0.928-0.966 | 0.814-0.831 | 0.928-0.966 | 0.815-0.832 | 0.924-0.962 |
| AJ888751 Austria (ALAD), AJ888752 | 0.804 | 0.940 | 0.805 | 0.940 | 0.803 | 0.940 | 0.804 | 0.936 |
| KC676609 Croatia (ALAD), KC676613 | 0.805-0.807 | 0.949-0.953 | 0.804-0.805 | 0.949-0.953 | 0.804-0.805 | 0.949-0.953 | 0.805-0.807 | 0.945-0.949 |
| FN377821 Hungary (ALAD), FN377822 | 0.810-0.807 | 0.957 | 0.808-0.811 | 0.957 | 0.805-0.808 | 0.957 | 0.810-0.807 | 0.953 |

| | | | | | | | | |
|---|---|---|---|---|---|---|---|---|
| AJ314600 Balkan (ALAD),<br>AJ314601 | 0.810-<br>0.807 | 0.949 | 0.808-<br>0.811 | 0.949 | 0.805-<br>0.808 | 0.949 | 0.810-<br>0.807 | 0.945 |
| GQ339483 Sweden (S-SCA),<br>Q339484, GQ339485, GQ339487, GQ339486, AJ223376 | 0.817-<br>0.841 | 0.974-<br>0.966 | 0.818-<br>0.842 | 0.974-<br>0.966 | 0.817-<br>0.841 | 0.974-<br>0.966 | 0.817-<br>0.841 | 0.970-<br>0.962 |
| AJ223368 Norway (S-SCA),<br>AJ223369 | 0.817-<br>0.818 | 0.962 | 0.817-<br>0.818 | 0.962 | 0.818-<br>0.819 | 0.962 | 0.817-<br>0.818 | 0.957 |
| EU439968 Germany (CE),<br>EU439969, EU439972, DQ016430, DQ016432, Y954722,<br>JN696358, KJ994776, JN696373, JN696372, JN696371,<br>JN696374, JN696376 | 0.805-<br>0.843 | 0.970-<br>0.957 | 0.804-<br>0.845 | 0.970-<br>0.957 | 0.807-<br>0.842 | 0.970-<br>0.957 | 0.805-<br>0.842 | 0.966-<br>0.953 |
| U22423 Belgium (CE),<br>J277075, AJ277076, AJ277034, AJ277033, AJ277032,<br>AJ277031, AJ277030 | 0.821-<br>0.797 | 0.957-<br>0.940 | 0.822-<br>0.798 | 0.957-<br>0.940 | 0.819-<br>0.796 | 0.957-<br>0.940 | 0.821-<br>0.797 | 0.953-<br>0.936 |
| AM695638 France (CE),<br>KT247592, KT247594, KT247595, KT247596, KT247597 | 0.825-<br>0.807 | 0.962-<br>0.957 | 0.824-<br>0.808 | 0.962-<br>0.957 | 0.824-<br>0.805 | 0.962-<br>0.957 | 0.825-<br>0.805 | 0.957-<br>0.953 |
| GQ339476 Sweden (N-SCA),<br>GQ339477, AM746297, AM746298, AM746310, AM746311,<br>AM746315, AM746316, AM746317, AM746318, AM746319,<br>GQ339478, GQ339480, GQ339482, GQ339473, GQ339474,<br>GQ339481, GQ339479, AM746320, AM746322, AM746324,<br>AM746325, AM746328, AM746329, AM746330, AJ223371,<br>AJ223374, AJ223375, AJ223380, Z48586, AM746332,<br>AM746331, AM746333, AY526219, U14137, AJ223377 | 0.828-<br>0.797 | 0.974-<br>0.949 | 0.829-<br>0.796 | 0.974-<br>0.949 | 0.828-<br>0.798 | 0.974-<br>0.949 | 0.828-<br>0.797 | 0.970-<br>0.945 |
| AJ238791 Denmark (DAN),<br>AJ278092, AJ278093 | 0.805-<br>0.786 | 0.953-<br>0.932 | 0.804-<br>0.787 | 0.953-<br>0.932 | 0.804-<br>0.784 | 0.953-<br>0.932 | 0.805-<br>0.786 | 0.949-<br>0.928 |

PUUV lineage affiliation is given at the end of sequence names: (ALAD) Alpe-Adrian lineage, (CE) Central European lineage, (DAN) Danish lineage, (FIN) Finnish lineage, (LAT) Latvian lineage, (N-SCA) North-Scandinavian lineage, (RUS) Russian lineage, (S-SCA) South-Scandinavian lineage. PUUV S segment sequences from colony bank voles 2, 4, 5, 8, 13, 14, 15, 17 and 18 were identical on nucleotide sequence level and therefore excluded.

| (b) Reference sequences of the M segment | PV798424<br>1gen14 Kraków<br>Poland (RUS) | | PV798425<br>4gen14 Kraków<br>Poland (RUS) | |
|---|---|---|---|---|
| | nt | aa | nt | aa |
| PV798424 1gen14 Kraków lab. colony Poland (RUS) | - | - | **0.998** | 1 |
| PV798425 4gen14 Kraków lab. colony Poland (RUS) | **0.998** | 1 | - | - |
| KX815397 KS14/118 Teleśnica Oszwarowa Poland (RUS) | 0.987 | 0.995 | 0.985 | 0.995 |
| KX815398 KS14/121 Teleśnica Oszwarowa Poland (RUS) | **0.998** | 1 | 0.996 | 1 |
| KF906513 KS13/855 Mikołajki Poland (LAT) | 0.804 | 0.922 | 0.805 | 0.922 |
| AF442614 Russia (RUS),<br>M29979, Z84205, AB433850, AB433851, AB433852 | 0.865-0.843 | 0.985-0.970 | 0.867-0.844 | 0.985-0.970 |
| Z70201 Finland (FIN), JQ319173, JQ319175, JQ319172, JQ319174, JN831951, NC_005223, HE801634 | 0.857-0.810 | 0.966-0.946 | 0.859-0.812 | 0.966-0.946 |
| KC676630 Croatia (ALAD),<br>KC676632, KC676634 | 0.826-0.823 | 0.922-0.917 | 0.828-0.825 | 0.922-0.917 |
| DQ518223 Germany (CE),<br>DQ518225, DQ518236, DQ518237, DQ518219, DQ518218, AJ238778, DQ518217, DQ518215, DQ518213, DQ518222, DQ518221, DQ518227, KJ994777 | 0.822-0.791 | 0.932-0.917 | 0.823-0.792 | 0.932-0.917 |
| U22418 Belgium (CE) | 0.802 | 0.927 | 0.804 | 0.927 |
| KT247603 France (CE), KT247602, KT247598, KT247599, KT247600, KT247601 | 0.813-0.796 | 0.927-0.893 | 0.815-0.797 | 0.927-0.893 |
| Z49214 Sweden (N-SCA),<br>AY526218, U14136 | 0.818-0.792 | 0.941-0.902 | 0.794-0.820 | 0.941-0.902 |

PUUV lineage affiliation is given at the end of sequence names: (ALAD) Alpe-Adrian lineage, (CE) Central European lineage, (FIN) Finnish lineage, (LAT) Latvian lineage, (N-SCA) North-Scandinavian lineage, (RUS) Russian lineage. PUUV M segment sequences from colony bank voles 2, 5, 6, 8, 9, 13, 14 and 17 were identical on nucleotide level and therefore excluded.

| (c) Reference sequences of the L segment | PV798426<br>1gen14 Kraków<br>Poland (RUS) | |
|---|---|---|
| | nt | aa |
| PV798426 1gen14 Kraków lab. colony Poland (RUS) | - | - |
| KX815400 KS14/118 Teleśnica Oszwarowa Poland (RUS) | 0.995 | 1 |
| KX815401 KS14/121 Teleśnica Oszwarowa Poland (RUS) | **0.997** | 1 |
| KX815399 KS13/855 Mikołajki Poland (LAT) | 0.866 | 0.978 |
| OL355107 Lithuania (LAT) | 0.868 | 1 |
| EF405801 Russia (RUS),<br>OP561837, OP561846, OP561849, OP561852 | 0.873-0.851 | 1-0.963 |
| JN831945 Finland (FIN),<br>HE801635, MT024595, MT024592 | 0.875-0.861 | 1-0.992 |
| MT514291 Germany (CE);<br>MT514293, MW656500, MN639749 | 0.868-0.851 | 0.985-0.956 |
| KT247604 France (CE),<br>KT247605, KT247606, KT247607, KT247608, KT247609 | 0.863-0.832 | 0.970-0.941 |
| AY526217 Sweden (N-SCA),<br>MN832779 | 0.873-0.854 | 0.978-0.970 |

PUUV lineage affiliation is given at the end of sequence names: (CE) Central European lineage, (FIN) Finnish lineage, (LAT) Latvian lineage, (N-SCA) North-Scandinavian lineage, (RUS) Russian lineage. On nucleotide level all PUUV L segment sequences from colony bank voles were identical, therefore only the sequence from individual 1 was included.
